# Dual role of CASP8AP2/FLASH in regulating epithelial-to-mesenchymal (EMT) plasticity

**DOI:** 10.1101/2023.03.12.532282

**Authors:** Madison Catalanotto, Joel Markus Vaz, Camille Abshire, Reneau Youngblood, Min Chu, Herbert Levine, Mohit Kumar Jolly, Ana-Maria Dragoi

**Affiliations:** Department of Molecular and Cellular Physiology, LSUHSC-Shreveport, Louisiana, USA; Feist-Weiller Cancer Center, INLET Core, LSUHSC-Shreveport, Louisiana, USA; LSU Health Shreveport, School of Medicine, Louisiana, USA; School of Biological Sciences, Georgia Institute of Technology, Atlanta, Georgia, USA; Research Core, LSUHSC-Shreveport, Louisiana, USA; Center for Theoretical Biological Physics, Northeastern University, Boston, MA, USA; Department of Physics, Northeastern University, Boston, MA, USA; Department of Bioengineering, Northeastern University, Boston, MA, USA; Centre for BioSystems Science and Engineering, Indian Institute of Science, Bangalore, India

**Keywords:** epithelial-to-mesenchymal transition (EMT), EMT transcription factors (EMT-TFs), caspase 8 associated protein 2 (CASP8AP2/FLASH)

## Abstract

Metastasis consists of sequential steps initiated by cancer cells invading from the primary tumor site into neighboring tissues, followed by entry into the circulatory system and completed by extravasation and growth in distal organs where secondary tumors are formed. Circulating tumor cells, thus, encounter and adapt to multiple environmental changes during their transition from the primary to the secondary tumor sites. Epithelial-to-mesenchymal transition (EMT) is a developmental program that consists of loss of epithelial features concomitant with acquisition of mesenchymal features. Activation of EMT in cancer facilitates acquisition of aggressive traits and cancer invasion. EMT plasticity (EMP), the dynamic transition between multiple hybrid states in which cancer cells display both epithelial and mesenchymal phenotypes, confers survival advantages for cancer cells in the constantly changing environment. Therefore, understanding the molecular mechanisms regulating intermediate phenotypic states along the E–M spectrum is critical. Core EMT transcription factors (EMT-TFs), ZEB, SNAI and TWIST families, play an important role in EMT and its plasticity. In the present study we characterize FLASH as a regulator of EMP and multiple EMT-TFs. We demonstrate that loss of FLASH gives rise to a hybrid E/M phenotype with high epithelial scores even in the presence of TGFβ, as determined by computational methods using expression of predetermined sets of epithelial and mesenchymal genes. We demonstrate that FLASH is regulating expression of multiple cell junction proteins with an established role in cancer progression and that its role in EMT is independent of its histone biogenesis role. Further, we show that FLASH expression in cancer lines is inversely correlated with the epithelial score, consistent with its function as a repressor of the epithelial phenotype. Nonetheless, activation of a distinct set of mesenchymal markers concomitant with epithelial markers reveals the complex role of FLASH in EMT and indicates that intermediate E/M states could arise from opposing control by FLASH on different families of EMT-TFs.

## Introduction

The most critical step in cancer pathogenesis, the development of metastasis, is responsible for the majority of cancer-associated deaths [1, 2]. Cancers with high and early rates of metastasis have a poor prognosis and are in general less sensitive to therapy. The physical translocation of cancer cells from the primary tumor site into the surrounding tissue and further to distant locations in the body followed by colonization of the secondary sites is mediated by activation of the epithelial-to-mesenchymal transition (EMT) [3-6]. A fundamental principle of EMT reprogramming is the acquisition of mesenchymal phenotype features by the epithelial tumor cells. During EMT reprogramming, a series of genetic and physiological changes induces loss of epithelial cell polarization. This is caused by transcriptional downregulation of multiple epithelial genes encoding cell-cell junction proteins required for the maintenance of basal-apical polarity, such as tight-junction proteins, E-cadherin, epithelial cellular adhesion molecule (EpCAM), desmoplakins, and cytokeratin [3, 4, 6], or by their post-translational modifications [7]. Concomitantly with the loss of epithelial markers, cancer cells increase the expression of mesenchymal markers such as N-cadherin, vimentin, fibronectins, and matrix metalloproteinases (MMPs) [3, 4]. Often, these EMT events are driven by one or more EMT transcription factors (EMT-TFs). The core EMT-TFs, include zinc-finger E-box-binding homeobox family (ZEB1 and ZEB2), SNAI family (SNAIL and SLUG), and Twist-related protein (Twist1 and Twist2) [8-10], are often co-expressed in various combinations and regulate each other to coordinate intricate EMT programs. Expression and activation of various EMT-TFs occur in response to multiple signaling pathways mediated by growth factors, cytokines and cues from the microenvironment such as TGFβ, EGF, FGF, PDGF, and fibronectin [11-16]. EMT-TFs pleiotropic functions are also responsible for cancer cells’ escape from apoptosis, acquisition of stemness properties, therapy resistance, and immune evasion [17-19].

EMT plasticity (EMP), a critical ability to adopt hybrid E/M features and transition between several intermediate EMT states, is believed to enable cancer cells to adjust to the changing environment during metastasis [20-24]. How gradual accumulation of mesenchymal markers is coupled or uncoupled from the concurrent loss of epithelial markers and the mechanisms regulating the dynamic switches between hybrid E/M states are still open questions. Nonetheless, these multiple intermediate phenotypic states along the E-M axis are believed to be fluid, interchangeable and likely facilitate invasion and resistance to chemotherapy [24-28]. Indeed, cancer cells in hybrid E/M states are enriched in populations of circulating tumor cells (CTCs) congruent with a role in survival in distinct environments and tissue dissemination [29-32].

FLASH/CASP8AP2 (caspase 8 - associated protein 2) was originally identified as a pro-apoptotic protein involved in Fas-mediated caspase-8 activation [33]. In acute lymphoblastic leukemia patients, loss of FLASH expression was correlated with poor treatment response and relapse [34, 35]. However, an anti-apoptotic role for FLASH has been described in multiple studies showing that FLASH can suppress apoptosis in both Fas-dependent and -independent manners [36-38]. Recently, a FLASH/AP-1 axis was identified in mediating lung cancer viability [39]. These studies suggest that FLASH can promote or inhibit apoptosis depending on the context and cell type involved. FLASH is also an important component of the complexes involved in the core histones precursor mRNA expression and processing [37, 40-42] together with NPAT (Nuclear protein of the ATM locus) [43] and SLBP (stem-loop binding protein) [44]. Loss of FLASH, thus, affects canonical histones pre-mRNA processing [37, 40], dampens core histones biogenesis and results in S-phase cell cycle arrest.

We have previously identified FLASH as a novel repressor of E-cadherin and other epithelial markers through a conserved mechanism involving ZEB1 protein degradation [38, 45]. Our studies showed that FLASH protects ZEB1 from proteasomal degradation while at the same time participates in SNAI EMT-TFs transcriptional regulation [38]. While loss of FLASH results in decreased ZEB1 expression, it also induces high levels of SNAIL and SLUG at both the mRNA and protein level in multiple cancer cell lines, suggesting a conserved mechanism of EMT-TFs regulation [38]. Despite expressing high levels of SNAIL and SLUG and even after TGFβ treatment, cells depleted for FLASH maintain high levels of E-cadherin, display a less invasive phenotype, and increased apoptosis in response to chemotherapy. Our initial studies uncovered that expression of several epithelial genes is repressed by FLASH, such as EpCAM and MARVELD3, most likely targets of ZEB1 [45-49]. However, other genes were controlled in a ZEB1-independent manner, highlighting the direct and indirect role of FLASH in transcriptional regulation [45]. These original studies support a role of FLASH as a repressor of the epithelial phenotype and a critical determinant of EMT.

Nonetheless, FLASH is required for core histones biogenesis and S-phase progression and thus, it could indirectly affect transcription because of chromatin architecture. S-phase cell cycle arrest alone, however, did not alter E-cadherin expression. This indicates that FLASH regulates E-cadherin independent of S-phase arrest caused by decreased histones biogenesis [38]. To what extent S-phase arrest influences the EMT landscape therefore remains an open question.

Here, we performed a comprehensive RNAseq analysis in pancreatic cancer cell line PANC-1 lacking three individual histone biogenesis and S-phase progression regulators - FLASH, NPAT and SLBP in order to distinguish between the role of FLASH in EMT-TFs regulation and its broader role in cell cycle progression. Recently, transcriptomics-based scoring, based on established epithelial and mesenchymal markers expression, was developed to quantify EMT phenotypes in cell lines and patients [50-54]. We employed EMT scoring methods to the RNAseq data and uncovered that FLASH plays a unique role in epithelial markers repression in cancer cells independent of its histone biogenesis function. In addition to its repression of epithelial markers, we also demonstrated that FLASH represses a distinct set of mesenchymal markers in cancer cells. The presence of both epithelial and mesenchymal markers in FLASH-depleted cells underscores its dual function in EMT plasticity, likely due to its unique opposing roles in regulation of ZEB and SNAI family of EMT-TFs. Finally, analysis of cancer gene expression data sets such as CCLE (Cancer Cell Line Encyclopedia) reveals a negative correlation between FLASH and the epithelial phenotype, as well as an inverse correlation with SNAIL and SLUG, once more highlighting its dual role in EMT. Our results, thus, reveal FLASH as a regulator of EMP in diverse cancer types and validate previous prediction models of EMT transition states [55, 56].

## Results

### E-cadherin regulation by FLASH is independent of histone biogenesis and S-phase arrest

We have previously shown that E-cadherin upregulation following depletion of FLASH was not the result of S-phase arrest but was most likely due to decreased ZEB1 levels [38]. Nonetheless, depletion of FLASH induced upregulation of multiple epithelial markers and several mesenchymal markers, some of which may be the result of pleiotropic genetic changes due to the S-phase arrest and changes in chromatin architecture. Thus, we sought to discriminate between S-phase arrest effects and a specific role of FLASH in EMT progression. For this, we performed single knockdowns (KD) of FLASH, NPAT, and SLBP, three core histones regulators, in PANC-1 pancreatic cancer cells. Loss of any one of these histone regulators [44] results in cell cycle arrest in S-phase (Fig. 1A). Importantly, individual depletion of FLASH, NPAT and SLBP did not affect expression of the other two regulators, suggesting that the subsequent transcriptional changes were specific (Fig. 1B). While depletion of any of these three factors was accompanied by low levels of histone H3 and histone H4, two of the nucleosome core components (Fig. 1C, Hist H3 and H4), depletion of FLASH triggered high expression of E-cadherin while depletion of SLBP had no effect and depletion of NPAT had a minimal effect (Fig. 1C, ECAD). Thus, loss of FLASH specifically, and not S-phase arrest nor loss of histones, causes upregulation of E-cadherin. These results confirmed our previous studies and showed that FLASH may play a unique role in epithelial phenotype regulation.

**Figure 1.**
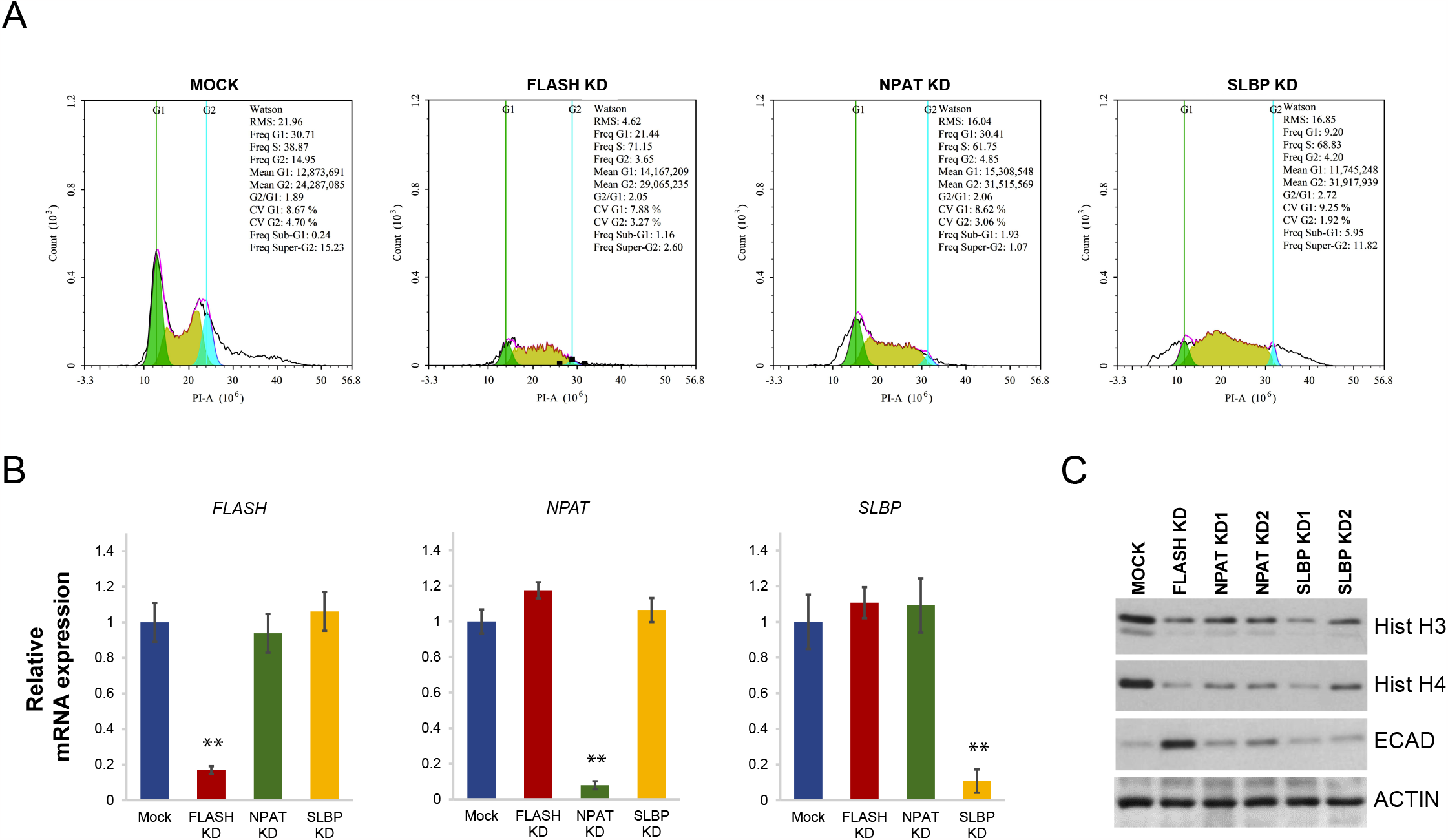
Depletion of histone biogenesis genes regulators triggers S-phase cell cycle arrest in cancer cells. (A) Cell cycle distributions in Mock-transfected (Mock) and FLASH (FLASH KD), NPAT (NPAT KD), and SLBP-depleted cells (SLBP KD) as measured by flow cytometry after PI staining. (B) *FLASH, NPAT* and *SLBP* gene expression in Mock-transfected and siRNA-transfected PANC-1 pancreatic cancer cells was evaluated by qPCR. The graphs represent the average of three independent experiments. The significance of differences was confirmed by Student t test for silencing efficiency (**, *p* < 0.01). (C) Core histones H3 (Hist H3), H4 (Hist H4) and E-cadherin (ECAD) protein levels were assessed by Western blot analysis in cells transfected with siRNA duplexes targeting FLASH (FLASH KD), NPAT (NPAT KD1 and NPAT KD2) and SLBP (SLBP KD1 and SLBP KD2). Actin was used as loading control.

### Genome-wide analysis of cancer cells reveals a specific role for FLASH as a transcriptional repressor

To understand further phenotypic similarities and differences after loss of FLASH, NPAT and SLBP in cancer cells, we profiled gene expression in Mock-transfected and siRNA-transfected PANC-1 cells at day 4 post-transfection. Differentially expressed genes (DEGs) in two independent experiments with an FDR < 0.1 were further used for functional analysis. Transcript levels of thousands of genes were altered in FLASH, NPAT and SLBP-depleted conditions (Fig. 2A and Table S1). Of those significant DEGs, approximately 1250 were commonly regulated by all three histone biogenesis factors (Table S2). Some DEGs were controlled by FLASH and NPAT only (1371 genes), FLASH and SLBP only (644 genes) or NPAT and SLBP only (1364 genes). Importantly, a large number of genes seem to be exclusively affected under the three siRNA-depleted conditions (Fig. 2A). Thus, 2403 genes were regulated only under FLASH-depleted condition (Table S3), 2378 genes under NPAT-depleted (Table S4) and 1092 under SLBP-depleted conditions (Table S5). This suggests that while certain genes are most likely affected because of S-phase arrest, others are not. Of all DEGs in the three siRNA conditions, only in FLASH-depleted cells were more genes upregulated than downregulated (Fig. 2B, 3308 genes upregulated *vs* 2359 genes downregulated, “All differentially expressed genes”). This difference was more striking when we analyzed genes specifically controlled by FLASH (1605 upregulated genes *vs* 798 downregulated), whereas NPAT or SLBP-depleted cells display a similar number of upregulated or downregulated genes (Fig. 2B, “Specific differentially expressed genes”). This is in agreement with the notion that FLASH acts as a transcriptional repressor.

**Figure 2.**
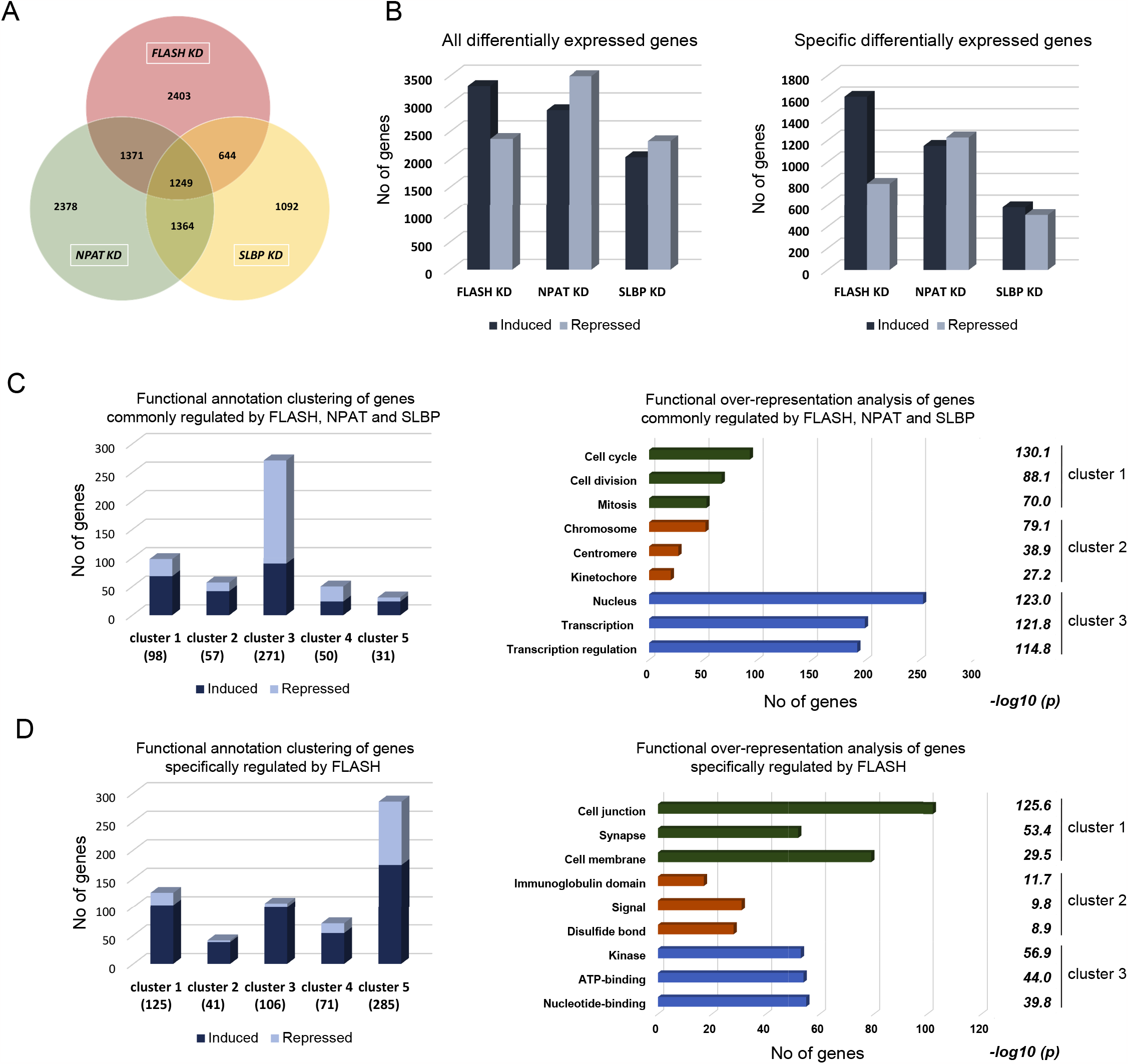
Transcriptome analysis of FLASH, NPAT and SLBP-depleted cancer cells reveals common and specific signatures. (A) Venn diagram showing the overlap among differentially expressed genes (DEGs) in PANC-1 cells depleted for FLASH, NPAT or SLBP (FLASH KD, NPAT KD and SLBP KD). (B) Number of DEGs between siRNA-transfected and Mock-transfected cells are shown for induced (dark blue) and repressed genes (light blue). Left panel shows all DEGs, right panel shows DEGs specifically regulated in FLASH, NPAT and SLBP-depleted cells. (C-D) Functional annotation clustering of DEGs commonly regulated by FLASH, NPAT and SLBP (C) or specifically regulated by FLASH (D). Left panel graphs indicates the top five enriched clusters and the number of DEGs in each cluster. Right panel graphs indicates functional over-representation in the top three clusters. The number of genes enriched in each term is shown on the x-axis. FDR value is shown for each enriched term.

Further, functional annotation clustering using Database for Annotation, Visualization and Integrated Discovery (DAVID) [57, 58] revealed that genes regulated by all factors are grouped in 15 cluster with an enrichment score >1.5 (Table S6), while genes specifically regulated by FLASH are grouped in 7 clusters with an enrichment score >1.5 (Table S7). The top 5 enriched clusters for commonly regulated and FLASH-specifically regulated genes are presented in Fig. 2C and 2D (left panels). Majority of DEGs specifically controlled by FLASH were upregulated in clusters 1-4 (Fig. 2D, left panel). Next, functional over-representation analysis of the 3 most enriched clusters was performed. The top functional categories for genes controlled by all three factors were related to “Cell cycle” and “Cell division” as expected (Fig. 2C, right panel, cluster 1). However, functional over-representation analysis for genes specifically controlled by FLASH revealed that most DEGs are enriched in “Cell junction” and “Cell membrane” categories (Fig. 2D, right panel, cluster 1). These results clearly delineate a unique role for FLASH in transcriptional control of genes involved in cell-cell junction, a role distinct from its role in histone biogenesis and S-phase progression.

### FLASH controls expression of cell junction genes deregulated in cancer

To confirm the role of FLASH in regulating genes associated with the epithelial phenotype, we further examined genes over-represented in the “Cell junction” category and performed a literature review to identify genes previously documented to play a role in cancer. We assembled a custom list of genes that function as suppressors or promoters of EMT and cancer progression (Table 1). Importantly, and consistent with the idea that FLASH is associated with loss of genes required for maintaining the epithelial phenotype, 21 genes previously identified as tumor and EMT suppressors were upregulated following FLASH depletion (Table 1, Upregulated). The role of MARVEL domain containing 3 (MARVELD3) protein, a tight junction associated protein, in inhibition of migration and EMT in hepatocellular carcinoma and pancreatic cancer has been described [48, 59]. The tumor suppressor CADM3/NECL1 (cell adhesion molecule 3) inhibits migration and invasion of glioma cells [60, 61] as well as tumorigenicity of colon cancer cells [62]. The tight junction-associated adaptor CGN (cingulin), inhibits tumorigenicity in mesothelioma and ovarian cancer [63, 64]. Epb41L3)/DAL-1 (erythrocyte membrane protein band 4.1 like 3) attenuates EMT in lung cancer [65-67], inhibits squamous cell carcinoma invasion [68, 69] and suppresses prostate cancer progression and metastasis [70, 71]. The tight junction protein CLDN6 (claudin-6) is a tumor suppressor in breast cancer [72-74] and colon carcinoma [75]. Thus, a significant set of genes responsible for epithelial junctions are downregulated by FLASH.

**Table 1:**
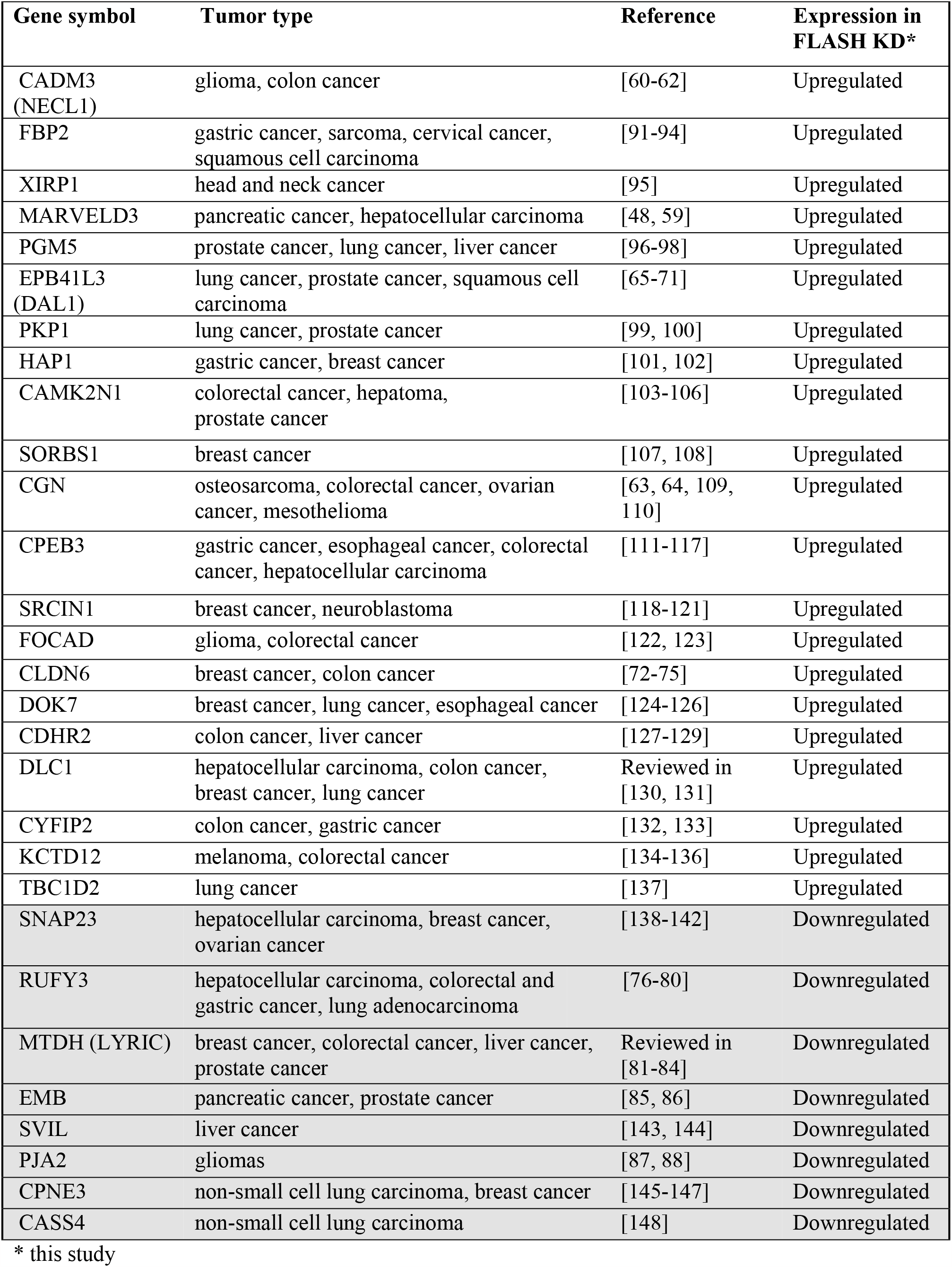
List of genes regulated by FLASH with a documented role in cancer

On the other hand, 8 genes with a described role in promoting EMT and cancer progression were repressed in FLASH-depleted cells (Table 1, Downregulated). For example, RUFY3 (RUN and FYVE domain containing 3), localizes to F-actin rich invadopodia, promotes EMT, invasion and metastasis in colorectal carcinoma, gastric cancer, and hepatocellular carcinoma [76-80]. Metadherin (MTDH/LYRIC/AEG-1) is involved in multiple signaling pathways such as PI3K/AKT, NF-κB, ERK, and Wnt/β-catenin and subsequently plays key roles in cancer progression, apoptosis evasion, invasion, and metastasis [81-84]. EMB (embigin) regulates cell motility and EMT in pancreatic cancer [85] and promotes prostate cancer progression [86]. PJA2 (praja ring finger ubiquitin ligase 2) is overexpressed in high-grade glioma tumors and attenuates Hippo signaling, thus, promoting tumor growth [87, 88]. Consistently, then, FLASH knockdown interferes with many aspects of the EMT program.

We validated markers in each category at the mRNA and protein level under four different conditions: FLASH KD, ZEB1 KD, NPAT KD and SLBP KD (Fig. 3A and 3D). This allowed us to confirm that epithelial markers such as MARVELD3 (Fig. 3B and 3D) is regulated by both FLASH and ZEB1 similarly to CDH [45], whereas CGN, CADM3 and CLDN6 (Fig. 3B and 3D) are specifically regulated by FLASH. Downregulation of EMB, LYRIC, RUFY3, and PJA2 (Fig 3C and 3D) is more specifically correlated with loss of FLASH (Figs. 3A-3D, FLASH KD). NPAT and SLBP do not play a significant role in regulation of majority of these markers (with the exception of RUFY3 and PJA2 which appear to be controlled post-translationally by SLBP) as evidenced by mRNA transcription and protein levels analysis (Figs. 3A-3D, NPAT KD and SLBP KD). Altogether, these results provide evidence that beyond E-cadherin regulation, FLASH plays a repressor role for several epithelial phenotype markers, either through ZEB1-dependent (e.g., CDH1 and MARVELD3) or ZEB1-independent mechanisms (e.g., CGN, CADM3 and CLDN6), and an activator role for junctional proteins with a negative role in EMT and cancer outcome. How FLASH mechanistically regulates transcription of some of these genes is still not understood. OVOL1, a transcription factor with a role in epithelial identity, is upregulated in FLASH-depleted cells (Fig. S1), possibly due to loss of ZEB1 and the release of their reciprocal inhibitory circuit [89, 90]. Thus, whether FLASH is a transcriptional repressor or activator itself, acts as a co-factor in transcription complexes or indirectly regulates transcription factors of the epithelial identity such as OVOL1, remains to be determined.

**Figure 3.**
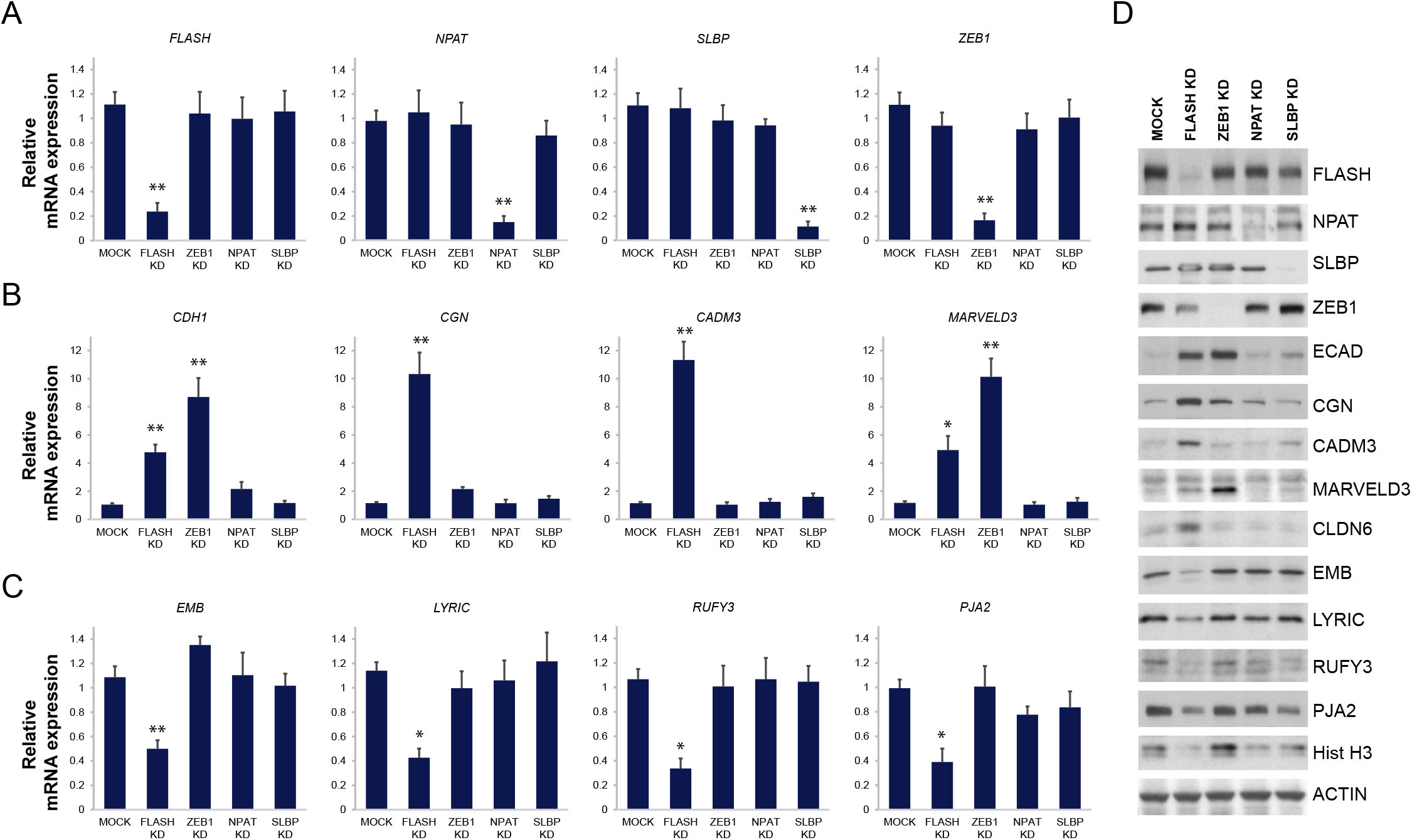
FLASH regulates expression of genes with roles in cell junction formation and EMT. (A-C) Gene expression profile in Mock, FLASH, NPAT, SLBP and ZEB1-depleted cells was evaluated by qPCR. (A) Graphs show mRNA levels of *FLASH, NPAT, SLBP* and *ZEB1*. (B-C) Four representative upregulated genes, *CDH1, CGN, CADM3* and *MARVELD3* (B) and, four representative downregulated genes, *EMB, LYRIC, RUFY3* and *PJA2* (C) are shown. The significance of differences was confirmed by Student t test for silencing efficiency (*, *p* < 0.05; **, *p* < 0.01). (D) Western blot analysis of genes tested by qPCR in A-C. Actin was used as loading control.

### Loss of FLASH drives a hybrid EMT phenotype with high epithelial scores

Recent studies have indicated that EMT can occur either completely or partially, the latter giving rise to cell states with mixed transcriptional and proteomic profiles. Furthermore, the detailed EMT status of cancer cells, as can be determined by an overall EMT “score”, can be a predictor of cancer progression, response to therapy and overall prognosis. Thus, we sought to evaluate the overall EMT score in cancer cells depleted for FLASH and the other histones regulators. For this we used two methods to calculate EMT scores: 76GS method (a 76-gene EMT signature developed using gene expression in non-small lung cancer cell lines) and KS method (the two-sample Kolmogorov-Smirnov test) [50-54]. The 76GS method considers 76 pre-determined genes and derives from the correlation coefficient between a particular gene expression and the expression of E-cadherin. Therefore, epithelial phenotypes score higher than mesenchymal phenotypes in the 76GS method. In the KS method, the scores vary on a scale from -1 to 1, the lower scores corresponding to the more epithelial phenotypes.

To gain insight into how these cells transition between intermediate phenotypes, we studied both baseline (untreated) cells and cells stimulated with TGFβ. TGFβ-induced EMT can occur at late stages of tumor development because of the activation of multiple signaling pathways [23, 149]. FLASH, NPAT and SLBP-depleted PANC-1 cells were treated with TGFβ for 48 hours. First, the 76GS and KS methods calculations showed a significant negative correlation in our samples analysis (Fig. 4A), consistent with previous validation [50]. Remarkably, both untreated and TGFβ-treated FLASH KD samples display the highest 76GS scores, with a shift from negative 76GS score values (mesenchymal) to positive score values (epithelial) (Fig. 4A, FLASH KD). NPAT-depleted cells also exhibited a more epithelial score, albeit, much lower than FLASH-depleted cells (Fig. 4A, NPAT KD). SLBP-depleted cells showed the least amount of change in the EMT scoring (Fig. 4A, SLBP KD). Interestingly, although TGFβ induced a more mesenchymal phenotype as evidenced by the lower 76GS and higher KS scores in all treated samples, FLASH-depleted cells retain a high epithelial score even after treatment (Fig. 4A, Mock TGFβ *vs* FLASH KD TGFβ). This is in agreement with our previous results, showing that E-cadherin expression in FLASH-depleted cells remains high despite TGFβ treatment [38]. Furthermore, *FLASH* and *ZEB1* expression showed a strong negative correlation with the 76GS score, whereas *CDH1* showed a positive correlation as expected (Fig. 4B, top panels). The same trend was observed when only expression of the 76GS epithelial genes was taken into account (Fig. 4B, lower panel).

**Figure 4.**
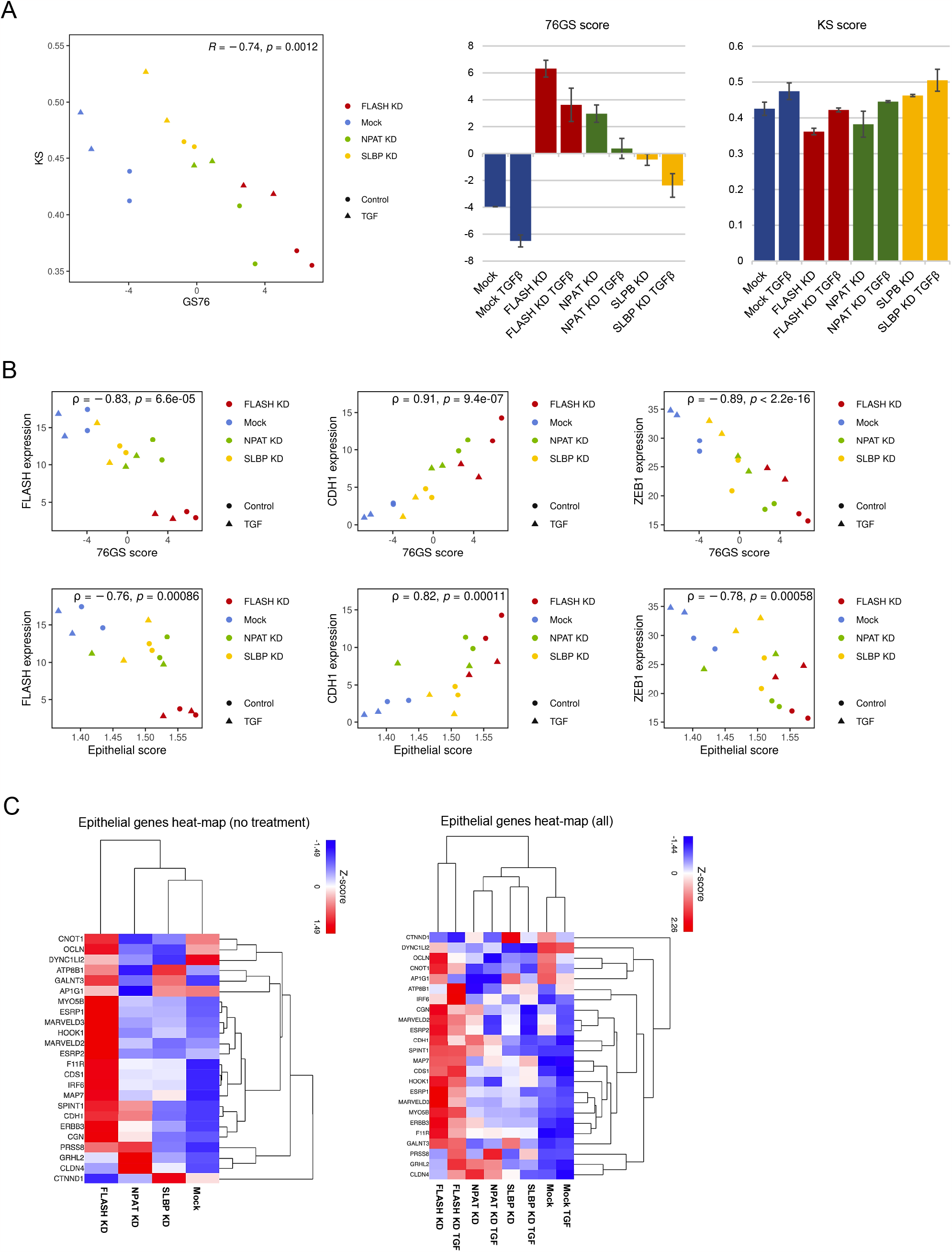
EMT phenotype quantification confirms the role of FLASH as an inhibitor of the epithelial phenotype. (A) Left panel: EMT scoring by 76GS and KS methods in TGFβ-treated and untreated Mock, FLASH, NPAT and SLBT-depleted cells. Two independent experiments were used for the analysis. KS and 76GS pairwise relation is assessed by Pearson’s correlation coefficient (*R*) and *p*-value (*p*). Right panels: bar plots showing average 76GS and KS EMT scores for individual conditions. (B) Scatter plots of *FLASH, CDH1* and *ZEB1* levels and the 76GS score (top panel) or the Epithelial score (lower panels) in TGFβ-treated and untreated Mock, FLASH, NPAT and SLBT-depleted cells. Spearman’s correlation coefficient (ρ) and corresponding p-value (*p*) are reported. Two independent experiments were used for the analysis. (C) Heat map analysis of epithelial genes [150] altered in FLASH, NPAT and SLBT-depleted cell under no treatment (left panel) and TGFβ-treated conditions (right panel).

Finally, we performed hierarchical clustering in our data groups for a subset of epithelial genes identified in a pan-cancer analysis from 11 tumor types as being associated with EMT and its clinical relevance [150] (Table S8). We observed that the majority of these genes were specifically and highly upregulated in FLASH-depleted cancer cells in comparison to Mock, NPAT and SLBP-depleted cells (Fig. 4C, No treatment). Their expression remains high even after TGFβ treatment, suggesting that FLASH-depleted cells may be refractory to TGFβ treatment (Fig. 4C, All). However, we cannot exclude the possibility that prolonged TGFβ treatment will cause a more pronounced effect on downregulation of epithelial markers as previously demonstrated [149]. Altogether, these results support the premise that FLASH acts as a repressor of critical aspects of the epithelial phenotype and substantiate the EMT scoring metrics as valuable tools for phenotype quantification.

### Loss of FLASH drives activation of certain mesenchymal genes signatures

Contrary to ZEB1 downregulation, SNAI family of EMT-TFs (SNAIL and SLUG) were shown to be upregulated in FLASH-depleted cells [38]. This suggests that FLASH plays dual and opposing roles in EMT-TFs regulation. SNAIL upregulation strongly correlated with loss of FLASH, but not cell cycle arrest (Fig. 5A). Importantly, SNAIL/SLUG regulation by FLASH is conserved in other cancer cell lines, such as cervical cancer and breast cancer (data not shown). Thus, a ZEB1^low^/SNAIL^high^/E-cadherin^high^ cell population emerges in FLASH-depleted cells (Fig. 5B, left panel). By contrast, ectopic expression of SNAIL in wild-type cells is sufficient to repress E-cadherin expression, similar to overexpression of FLASH or ZEB1 alone (Fig. 5B, right panel). Altogether these results suggest that FLASH may be required for SNAIL repressor functions during EMT. Nonetheless, in addition to their transcriptional repressor roles, EMT-TFs can activate expression of other EMT-TFs [151] as well as mesenchymal markers such as metalloproteases and collagens [152, 153].

**Figure 5.**
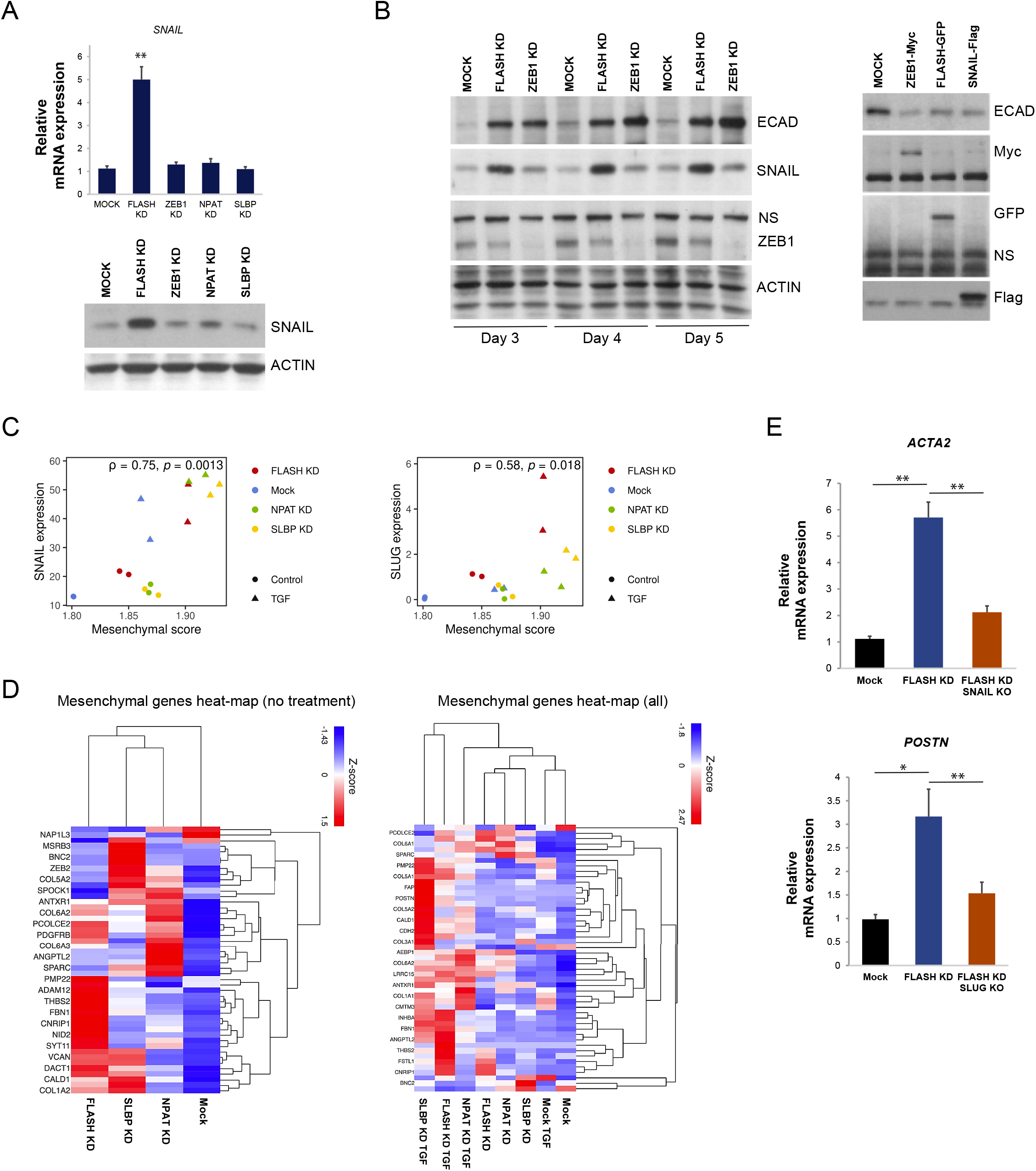
Loss of FLASH causes a hybrid EMT phenotype with a distinctive mesenchymal genes signature. (A) *SNAIL* EMT-TF upregulation in FLASH KD cells was determined by qPCR at mRNA level and Western blot analysis at protein level. (B) Hybrid EMT phenotype in FLASH-depleted cells as evidenced by distinct expression patterns of ECAD, SNAIL and ZEB1 (left panel). Overexpression of either ZEB1 (ZEB1-Myc), FLASH (FLASH-GFP) or SNAIL (SNAIL-Flag) inhibits E-cadherin expression (ECAD) in cancer cells (right panel). (C) Scatter plots of *SNAIL* and *SLUG* levels and the Mesenchymal score in TGFβ-treated and untreated FLASH, NPAT and SLBT-depleted cells. Spearman’s correlation coefficient (ρ) and corresponding p-value (*p*) are reported. Two independent experiments were used for the analysis. (D) Heat map analysis of mesenchymal genes [150] altered in FLASH, NPAT and SLBT-depleted cell under no treatment (left panel) and TGFβ-treated conditions (right panel). (E) Mesenchymal markers *ACTA2* and *POSTN* expression was determined by qPCR at mRNA level in WT, SNAIL KO and SLUG KO in control cells and FLASH-depleted cells. The significance of differences was confirmed by Student t test for silencing efficiency (*, *p* < 0.05; **, *p* < 0.01).

We next investigated if mesenchymal and epithelial markers coexist in cancer cells under our experimental conditions. First, we observed a significant positive correlation between expression of *SNAIL* and *SLUG* and the samples’ mesenchymal score (Fig. 5C) and hallmark EMT score (Fig. S2). Second, hierarchical clustering of mesenchymal genes with an identified role in EMT [150] (Table S9) showed that transcript levels of distinct sets of mesenchymal genes are specifically upregulated under the three conditions (Fig. 5D, No treatment). This was also true for TGFβ treatment (Fig. 5D, All). We hypothesized that SNAIL and SLUG upregulation in FLASH-depleted cells is at least partially responsible for the higher mesenchymal score. To validate this premise, we generated individual SNAIL and SLUG knock-out lines (Fig. S3, SNAIL KO and SLUG KO). Indeed, FLASH depletion in SNAIL KO or SLUG KO cells (FLASH KD/SNAIL KO and FLASH KD/SLUG KO) showed that upregulation of canonical mesenchymal markers such as alpha-smooth muscle actin and periostin is reversed by loss of SNAIL or SLUG (Fig. 5E, *ACTA2* and *POSTN*). These results provide evidence that although FLASH appears to play a crucial role in inhibiting the epithelial phenotype (Fig. 4), potentially promoting destabilization of the epithelial cells junctions, it may promote generation of hybrid E/M states by also inhibiting SNAI EMT-TFs expression.

### FLASH expression correlation with EMT markers in different cancer cell lines

To substantiate our findings regarding the role of FLASH in EMT plasticity, we analyzed the Cancer Cell Line Encyclopedia (CCLE) gene expression datasets. First, we observed that the 76GS EMT score negatively correlated with *FLASH* expression (Fig. 6A, FLASH expression/76GS) across a number of cancers. Given that 76GS score is higher for epithelial cells, this suggests that in multiple cancer lines FLASH acts as an epithelial phenotype repressor. A more in-depth tissue specific analysis revealed that certain cancer types such as liver and bone exhibit a very strong negative correlation (ρ < - 0.5), whereas lung cancer exhibited a strong negative correlation (ρ < - 0.2) between 76GS score and *FLASH* expression (Fig. 6B, CCLE: 76GS scores by cancer). As expected, a negative correlation was identified between *CDH1* and *FLASH* (Fig. 6A, CDH1 expression/FLASH expression). Notably, bone and liver cancer displayed the most significant negative correlation (Fig. 6B, CCLE: CDH1 *vs* FLASH), suggesting that FLASH may play a major role in EMT in these cancer types.

**Figure 6.**
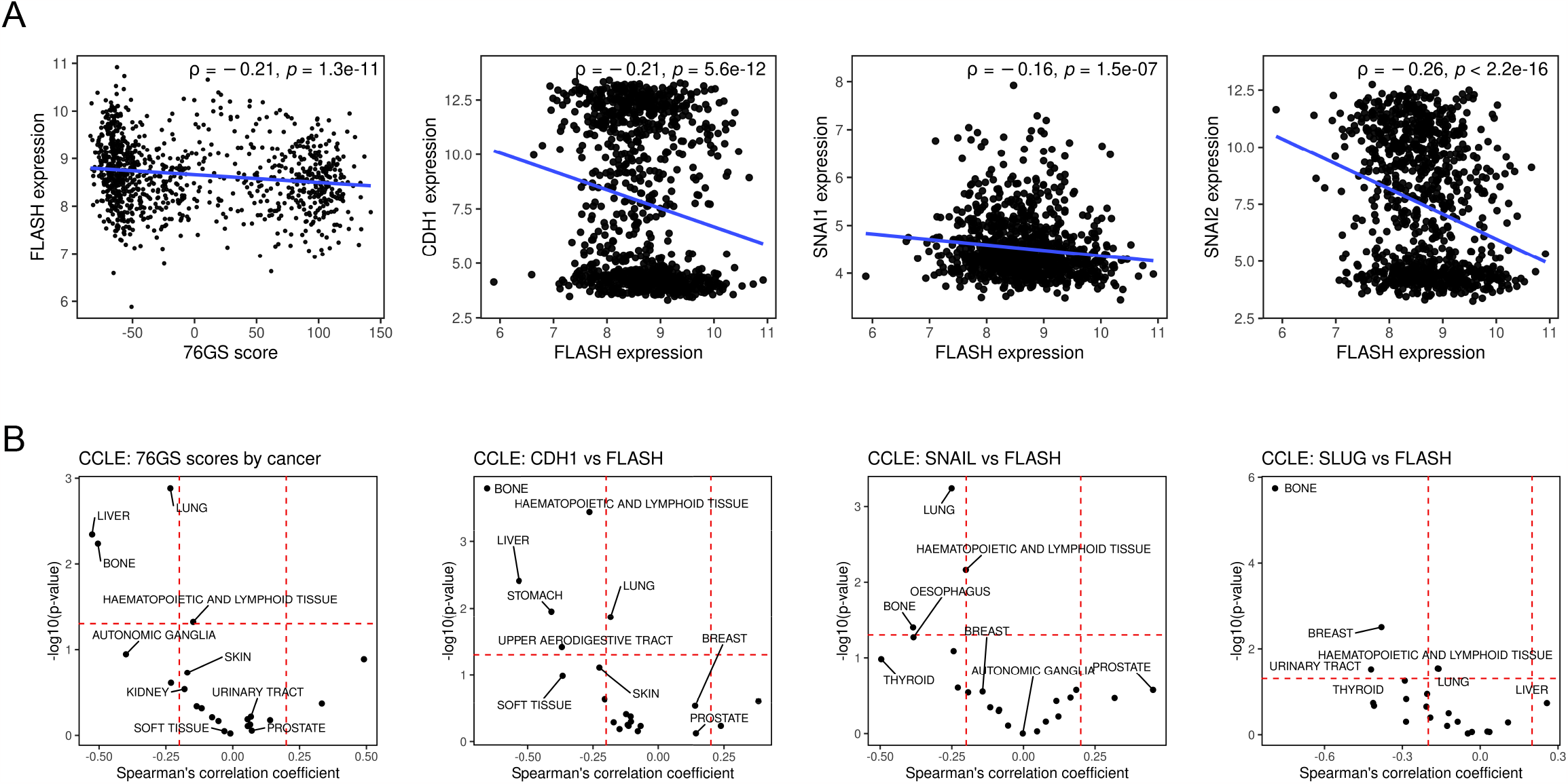
Correlation of FLASH expression with epithelial and mesenchymal markers in diverse cancer lines. (A) Scatter plots showing correlations between *FLASH* expression and 76GC score, *FLASH* and *CDH1* expression, *FLASH* and *SNAIL* expression, *FLASH* and *SLUG* expression in cell lines from CCLE. Spearman’s correlation coefficient (ρ) and corresponding p-value (*p*) are reported. (B) Tissue specific grouping of CCLE cells lines reveals tissues with a strong significant correlation (ρ > 0.2 and *p* value < 0.01).

Finally, we analyzed expression of *FLASH* and the *SNAI* family of EMT-TFs in the CCLE collection. Importantly, a negative correlation between both *SNAI* family members and *FLASH* expression was identified (Fig. 6A, SNAIL expression/FLASH expression and SLUG expression/FLASH expression). Similarly, strong negative correlation was identified between *SNAIL/SLUG* and *FLASH* in bone cancer types but also lung and breast cancer (Fig. 6B, CCLE: SNAIL *vs* FLASH and CCLE: SLUG *vs* FLASH). These inverse correlations of *FLASH* with both epithelial markers (*CDH1*) and EMT-TFs (*SNAIL* and *SLUG*) in bone cancers confirm its complex dual repressor role in EMT. In addition to *SNAIL* and *SLUG*, vimentin (*VIM)*, a mesenchymal marker abundantly expressed in sarcomas also negatively correlates with *FLASH* in bone cancers (Fig. S4). Differently from epithelial cancers, sarcomas have mesenchymal origins, although some sarcomas can display “epithelial-like” features [154]. Interestingly, a phenotypic switch with the acquisition of epithelial markers is beneficial for the clinical outcome in sarcoma patients [155]. Whether the opposing mechanisms of FLASH control over ZEB1 and SNAIL/SLUG are conserved in sarcomas, or are selectively activated, remains to be determined.

## Discussion

The major challenges in understanding EMP arise from the complexity of EMT programs and the multiplicity of mechanisms involved in generating hybrid EMT states. Phenotypic plasticity may confer advantages during migration and invasion together with chemotherapy resistance [24-28], yet, we do not fully understand how transitional states are generated or maintained. In general, EMT programs are executed by core EMT-TFs. Co-regulators of core EMT-TFs and multiple microRNAs together with long non-coding RNAs also contribute to activation of EMT. For the cells to progress completely through EMT, transcriptional changes in both epithelial and mesenchymal markers must be triggered, and genes are either repressed (epithelial markers) or activated (mesenchymal markers). However, in hybrid EMT states, epithelial and mesenchymal markers coexist, suggesting that either distinct EMT-TFs control repression of epithelial genes and activation of mesenchymal genes or that the repressor/activator functions of the same EMT-TFs are decoupled in these EMT states.

Here we describe a hybrid EMT phenotype generated by loss of FLASH in pancreatic cancer cells. Cells depleted for FLASH acquire a ZEB1^low^/E-cadherin^high^ phenotype. Multiple genes involved in cell-cell junction formation are similarly specifically upregulated in FLASH-depleted cells (Fig. 2D and Fig. 3), suggesting that FLASH acts as a repressor of the epithelial phenotype. Because loss of FLASH is associated with decreased ZEB1 expression, we hypothesize that some of the epithelial markers are de-repressed when this core EMT-TF is lost, such as CDH1 and MARVELD3 (Fig. 3). However, other epithelial markers are specifically regulated by FLASH in a ZEB1-independent manner (Fig. 3, CGN, CADM3 and CLDN6). This supports the idea that FLASH depletion is releasing transcriptional repression for other epithelial markers through a different mechanism. One possibility is that loss of ZEB1 in FLASH-depleted cells triggers expression of epithelial lineage transcription factors such as OVOL1 (Fig. S1). This implies that the increase in OVOL1 expression, controlled by the ZEB1:OVOL1 ratio and their mutual inhibition [89, 90] could promote epithelial cellular identity. Adding complexity to this mechanism is the fact that other cell membrane proteins with a role in promoting EMT and cancer progression are downregulated in FLASH-depleted cells (Fig. 3, EMB, LYRIC, RUFY3 and PJA2). We cannot exclude the possibility that FLASH itself is a transcriptional repressor/activator for some of these epithelial genes as FLASH has been shown previously to function as a transcriptional co-factor [156, 157].

EMT scoring using preset lists of genes with an established role as either epithelial or mesenchymal markers strongly associated loss of FLASH with a more epithelial phenotype. The high epithelial score is supported by the heatmap analysis of epithelial genes that underlines the role of FLASH but not NPAT or SLBP in suppressing most of these genes (Fig. 4C). Even so, analysis of mesenchymal gene expression identified distinct sets of genes being upregulated when either FLASH, NPAT or SLBP are depleted in cancer cells (Fig. 5D). We hypothesize that upregulation of many mesenchymal genes in FLASH-deleted cells may be the result of SNAI EMT-TFs high expression (Fig. 5A-B). Consistent with this premise, SNAIL and SLUG KO cells show a decrease in gene expression for *ACTA2* and *POSTN*, two mesenchymal markers upregulated in FLASH-depleted cells. Interestingly, as observed, knockout of a single SNAI family member is insufficient to block completely the expression of *ACTA2* or *POSTN*. This is mostly likely due to their redundant functions and the fact that FLASH depletion causes high expression of both SNAIL and SLUG [38]. Experiments targeting both SNAIL and SLUG to assess mesenchymal genes controlled by SNAI family in FLASH-depleted cells are in progress.

Nonetheless, certain mesenchymal genes are activated in NPAT and SLBP-depleted cells. How NPAT and SLBP are controlling expression of these sets of mesenchymal markers is unknown. Because of a certain degree of specificity, we suspect that different pathways and possibly EMT-TFs other than the core factors may be involved.

The ZEB1^low^/SNAIL^high^/E-cadherin^high^ phenotype described in our studies is therefore driving a hybrid E/M phenotype in which epithelial and mesenchymal markers coexist. This is significant for several reasons. Our data experimentally supports previous computational models showing that with variations in *ZEB1* and *SNAIL* expression levels cells occupy multiple mono or bistable states [55]. Nonetheless, these cells are susceptible to chemotherapy and are less invasive as we previously described [38]. Thus, expression of multiple epithelial markers may override expression of mesenchymal markers and force the hybrid E/M phenotype into a less invasive and therapy sensitive state. The question becomes, which epithelial markers or which combination of epithelial makers are necessary to maintain the hybrid E/M state in a less aggressive phenotype.

Given that neither loss of ZEB1, nor NPAT or SLBP causes upregulation of SNAI, we infer that FLASH plays a direct role in SNAI EMT-TFs transcriptional regulation. Indeed, gene expression analysis in CCLE collection shows that *FLASH* expression inversely correlated with both *SNAIL* and *SLUG* (Fig. 6). An important finding in our studies is that overexpression of SNAIL in wild-type PANC-1 cells suppresses E-cadherin expression (Fig. 5B, right panel) while the same is not true in FLASH-depleted cells (Fig. 5B, left panel). Combined with the finding that upregulation of several mesenchymal markers is SNAIL/SLUG-dependent, we propose that the “repressor” functions of SNAIL and SLUG are inhibited, while the “activator” functions are maintained. Several considerations could explain this: (i) loss of FLASH affects SNAI binding to its targets’ promoters or the formation of repressor complexes; (ii) SNAI “repressor” and “activator” functions are decoupled due to preferential redistribution in transcriptional complexes; (iii) upregulation of epithelial lineage transcription factors such as OVOL1 overrides the EMT-TFs repressor functions; (iv) multiple fully activated pathways are required for epithelial markers repression. First, very little is known about the transcriptional activity of FLASH itself, its direct targets and potential partition in transcriptional complexes, thus a direct role of FLASH in repression of epithelial markers cannot be excluded. Similarly, loss of FLASH could alter repressor complexes, such as the SIN3A complex (SIN3A/HDAC), which together with SNAIL is recruited to the *CDH1* promoter [158]. Second, while the role of EMT-TFs as repressors have been extensively investigated, their transcriptional activator roles are still unclear. ZEB1 has been shown to act not only as a major repressor of epithelial genes but also as an activator of mesenchymal genes when in complex with YAP1 [159, 160]. Less is known about the “activator” functions of SNAIL/SLUG, although several mesenchymal targets such as fibronectin, vimentin, SMA, MMP9, COX2, COL1A1, LEF have been identified [152]. ZEB1 itself is transcriptionally upregulated by SNAIL in response to TGFβ [151, 161]. Therefore, functional separation of EMT-TFs roles and preferential recruitment in either “repressor” or “activator” complexes could explain the generation of hybrid E/M phenotypes. Third, OVOL TFs overexpression (Fig. S1) could drive the epithelial phenotype, which is consistent with OVOL acting as a “molecular brake on EMT” [162]. Finally, it is possible that, with loss of ZEB1, FLASH-depleted cells do not reach the tipping point to irreversibly generate an aggressive hybrid E/M phenotype despite high levels of SNAIL and SLUG and expression of multiple mesenchymal markers.

All these scenarios could explain the distinctive hybrid E/M phenotype generated by loss of FLASH in cancer cells and generate new avenues of investigation into the repressor and activator functions of EMT-TFs. Further studies should shed light on the complex role of FLASH in EMT and its potential as a target for cancer therapy.

## MATERIALS AND METHODS

### Cell culture conditions

PANC-1 (CRL-1469) cells were obtained from ATCC and grown in DMEM media (Genesee) supplemented with 10% FBS. Cells were cultured at 37°C in a 5% CO2 incubator. The cell lines was authenticated by short tandem repeat (STR) profiling and tested for *Mycoplasma* contamination.

### RNAi assays

Cells were reverse transfected with Dharmafect1 (Dharmacon) and a pool of two individual siRNA silencing duplexes (25nmol/L each, 50 nmol/L total). All siRNA duplexes were purchased from Horizon/Dharmacon: FLASH (D-012413-01-0005 and D-012413-17-0005), NPAT (D-019599-02-0005 and D-019599-19-0005), SLBP (D-012286-01-0005 and D-012286-02-0005), ZEB1 (D-006564-02-0005 and D-006564-03-0005). Duplexes used in this study were validated for KD efficiency among 4 individual duplexes targeting the same gene. The siRNA transfection was allowed to proceed for 48h before treatment with TGFβ (100ng/ml) or control media was added for another 48h.

### RNA sequencing

RNA-seq was carried out in duplicates for all conditions. RNA was extracted from cancer cells using the RNeasy Plus Kit from Qiagen. Total RNA integrity was assessed on Agilent TapeStation 2200. Libraries were prepared using Illumina’s TruSeq Stranded mRNA kit. Libraries were analyzed on an Agilent TapeStation 2200 D1000 assay to determine average size and were quantitated using the Quanta PerfeCta NGS qPCR Quantification Kit. Libraries were normalized to 4nM, pooled, denatured, and diluted to approximately 1.8 pM. A 1% library of 1.8 pM PhiX was spiked in as an internal control. The library pool was sequenced on an Illumina NextSeq 550, with a read length of 2 × 75 base pairs. Two runs were performed. Base calling and quality scoring were performed with Illumina Real Time Analysis software (RTA). <u>Analysis</u>: Reads were aligned to the human genome (GRCh38.p10) using STAR_2.4.2a and counted using RSEM 1.2.31. Differentially expressed genes were identified with EBSeq 1.12. and filtered with a 0.1 FDR cutoff. Partek Flow was used for hierarchical clustering and heatmap construction. Genes with >1 transcripts per million (TPM) in at least 1 of 8 conditions (untreated and TGFβ-treated) were included in the analysis. Average of two independent experiments were used TPM analysis.

### Gene functional analysis

Gene Ontology (GO) analysis associated with the DEGs was performed using the Database for Annotation, Visualization and Integrated Discovery (DAVID), with higher enrichment score signifying more functional enrichment [57, 58] (https://david.ncifcrf.gov). Default values were used for functional annotation (Count: 2, EASE: 0.1).

### RNA extraction and quantitative PCR

RNA was extracted from cancer cells using the RNeasy Plus Kit from Qiagen. Total RNA and first-strand cDNA synthesis was performed using TaqMan Gene Expression Cells-To-Ct Kit (ThermoFisher) as previously described [38, 45]. mRNA levels were determined by quantitative real-time PCR using the Universal ProbeLibrary (Roche, Life Science) and LightCycler 480 Probes Master (Roche, Life Science). For the LightCycler 480 Probes Master the thermal cycling was carried out using a LightCycler 96 instrument (Roche Diagnostics) under the following conditions: 95°C for 5 minutes and 40 cycles at 95°C for 10s and 60°C for 25s. Relative quantification was performed using 2^-ΔΔCT^ method. Gene expression was normalized to GAPDH as reference gene. A complete list of primers and probes used in the study are presented in Table S10. All experiments were performed at least three times. Data are presented as average of three or four repeats. The analysis utilized Student’s *t* tests to determine significance. Values of *P* < 0.05 were considered significant, values of *P* < 0.01 were considered highly significant.

### Western blot analysis

PANC-1 cells were lysed in RIPA buffer prior to SDS-PAGE analysis and immunoblotting. The primary antibodies used were E-cadherin (Clone 36; BD Transduction Laboratories), Actin (C-2; Santa Cruz), NPAT (sc-136007; Santa Cruz), SLBP (ab181972; Abcam). The following antibodies were all purchased from Cell Signaling: FLASH (D3T8Q), ZEB1 (E2G6Y), SNAIL (C15D3), SLUG (C19G7), Claudin-6 (E7U20), LYRIC (D5Y8R), RUFY3 (61460S), PJA2 (40180S), GFP (D5.1), Myc (9B11), Histone H3 (D1H2) and Histone H4 (D2X4V). The following antibodies were all purchased from Sigma-Aldrich: MARVELD3 (AV44715), CGN (HPA027657), CADM3 (SAB1411161), EMB (SAB2700691) and Flag (F3165). Secondary antibodies horseradish peroxidase (HRP)-anti-mouse and anti-rabbit (1:5,000) were from Jackson Laboratories.

### Generation of PANC-1 stable cell lines

SNAIL and SLUG knockout PANC-1 cell lines were generated using lentiCas9-Blast (Addgene plasmid 52962) and lentiGuide-Puro (Addgene plasmid 52963) as previously described [163]. Guide RNAs used in this study are listed in Table S11.

### Cell cycle analysis

Cancer cells transfected with siRNA and control cells were collected after 72h, fixed in 70% cold ethanol and incubated for 30 minutes in a 1xPBS solution with 0.1% Triton X-100, 100µg/ml RNA-ase and 50µg/ml Propidium iodide (PI). Cell were analyzed on Novocyte Quanteon (Agilent).

### EMT score calculation methods

The raw counts obtained from the samples were normalized to their transcripts per million (TPM) values and log2 transformed before their EMT scores were calculated. 76GS and KS scores were calculated using algorithms standardized from microarray scoring metrics to fit RNA-seq data [164]. 76GS scoring metric, derived from non-small cell lung cancer (NSCLC), uses 76 genes to calculate the scores of the samples based on the correlation coefficient of that gene with the *CDH1* expression [54]. A higher 76GS score represents a more epithelial sample. The two-sample Kolmogorov–Smirnov test (KS score) calculates scores based on several epithelial and mesenchymal genes. A higher KS score represents the sample’s mesenchymal nature [53]. Single-sample Gene Set Enrichment Analysis (ssGSEA) [165] was performed using GSEAPY in Python, and the epithelial and mesenchymal cell line signatures were derived from the KS scoring genes, while the EMT signature was taken from the hallmark gene sets in the Molecular signatures database (MSigDB) [166].

### Pan-cancer samples analyses

The Cancer Cell Line Encyclopedia (CCLE) microarray dataset [167] was downloaded from the CCLE website (https://sites.broadinstitute.org/ccle/). Samples were separated according to the cancer type, and the axes were correlated using Spearman’s correlation. A value of ρ > 0.2 was considered significantly correlated.

## Statistical analysis

Two-tailed two-sample Student’s t-test was calculated between samples to check for a significant increase or decrease in their respective scores. A *p*-value (labeled across samples in boxplots) of < 0.05 was considered significant. R version 4.1.2 and Python 3.9.7 were used for all analysis, and package “ggplot2” in R was used for plotting functions.

## Data availability statement

All RNAseq data will be deposited in GEO repository and the accession number made available upon publication.

## Author Contributions

AMD conceptualized the project and acquired funds for the project. MC, JMV, CA, RY, MC and AMD performed the experimental work and data analysis. HL and MKJ oversaw the computational analysis for the paper. AMD wrote the original draft. AMD, HL, MKJ and MC reviewed and edited the final manuscript.

## ACKNOWLEDGMENTS

The authors would like to acknowledge the Research Core facility at LSUHSC-Shreveport that performed the RNAseq experiment and provided the list of DEGs. This work was supported by start-up funds from LSUHSC-Shreveport and Feist-Weiller Cancer Center Shreveport to AMD. Additional funding: P20GM134974 from National Institute of General Medical Sciences of the National Institutes of Health to AMD. MKJ was supported by Ramanujan Fellowship (SB/S2/RJN-049/2018) awarded by SERB, DST, Government of India. HL was supported by the National Science Foundation grant PHY-2019745.

