## Supplemental Figure Legends for "Dual role of CASP8AP2/FLASH in regulating epithelial-to-mesenchymal (EMT) plasticity"

**Supplementary Information**

**Table S1**
List of DEGs in Mock *vs* FLASH KD, Mock *vs* NPAT KD and Mock *vs* SLBP KD. Values of PostFC >1 are upregulated genes, values <1 are downregulated genes. Files shows DEGs identified in two independent experiments with a 0.1 FDR cutoff.

**Table S2**
List of commonly regulated DEGs in Mock *vs* FLASH KD, Mock *vs* NPAT KD and Mock *vs* SLBP KD. Values of PostFC >1 are upregulated genes, values <1 are downregulated genes. Files shows DEGs identified in two independent experiments with a 0.1 FDR cutoff.

**Table S3**
List of DEGs specifically regulated in FLASH KD. Values of PostFC >1 are upregulated genes, values <1 are downregulated genes. Files shows DEGs identified in two independent experiments with a 0.1 FDR cutoff.

**Table S4**
List of DEGs specifically regulated in NPAT KD. Values of PostFC >1 are upregulated genes, values <1 are downregulated genes. Files shows DEGs identified in two independent experiments with a 0.1 FDR cutoff.

**Table S5**
List of DEGs specifically regulated in SLBP KD. Values of PostFC >1 are upregulated genes, values <1 are downregulated genes. Files shows DEGs identified in two independent experiments with a 0.1 FDR cutoff.

**Table S6**
Functional clustering of commonly regulated DEGs in Mock *vs* FLASH KD, Mock *vs* NPAT KD and Mock *vs* SLBP KD.

**Table S7**
Functional clustering of DEGs specifically regulated in FLASH KD.

**Table S8**
TPM values for epithelial genes used for heatmap analysis in Partek Flow.

**Table S9**
TPM values for mesenchymal genes used for heatmap analysis in Partek Flow.

**Table S10**
List of qPCR primers and corresponding probe number from Universal Probe Library (Roche).

**Table S11**
Guide RNAs used to perform knockout (KO) experiments for SNAIL and SLUG.

**Figure S1**
OVOL1 gene expression in Mock and FLASH KD was quantified by qPCR in three independent experiments. The significance of differences was determined by Student *t* test (**, $p<0.01$).

**Figure S2**
Correlation of SNAIL and SLUG expression with Hallmark EMT score in Mock, FLASH KD, NPAT KD and SLBP KD conditions.

**Figure S3**
WT, SNAIL KO and SLUG KO pancreatic cancer cells PANC-1 were Mock-transfected (MOCK) or siRNA transfected with duplexes targeting FLASH (FLASH KD). Western blot was perform to show induction of SNAIL and SLUG in FLASH-depleted cells in WT cells and lack of proteins in SNAIL and SLUG KO cells.

**Figure S4**
Tissue specific grouping of CCLE cell lines showing correlation between FLASH and Vimentin expression.
