## Supplementary figures and images for "Dual role of CASP8AP2/FLASH in regulating epithelial-to-mesenchymal (EMT) plasticity"

### Supplemental Figure 1

Supplemental Figure 1

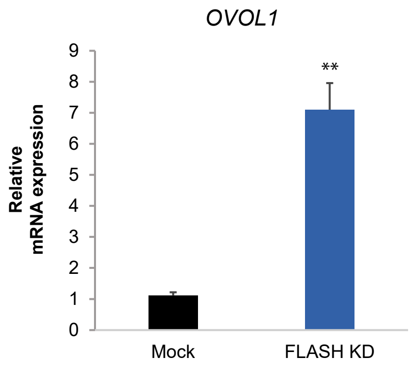

### Supplemental Figure 2

#### Supplemental Figure 2

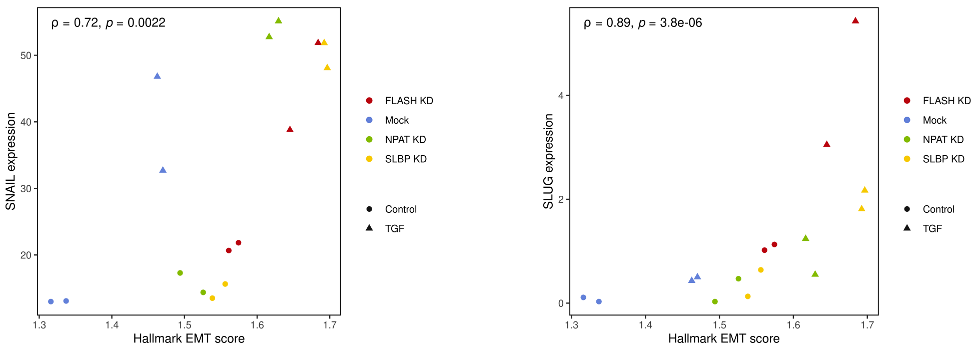

### Supplemental Figure 3

Supplemental Figure 3

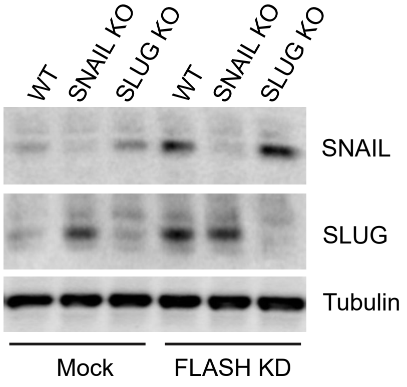

### Supplemental Figure 4

#### Supplemental Figure 4

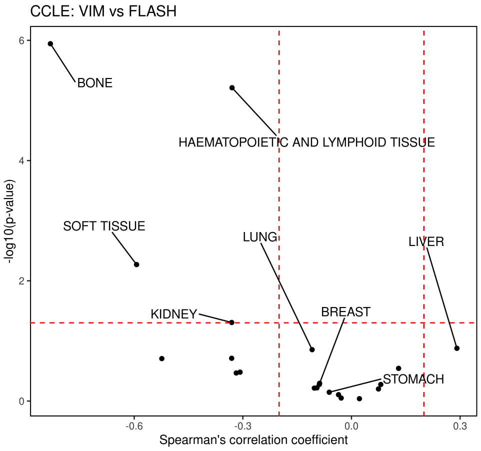
